# Mechanism of selectivity reveals novel antifolate drug interactions

**DOI:** 10.1101/2020.09.18.304022

**Authors:** Shannon Lynn Kordus, Elise A. Lamont, Michael D. Howe, Allison A. Bauman, William McCue, Barry Finzel, Anthony D. Baughn

## Abstract

Antimicrobial agents that target a specific pathogen of interest is the gold standard in drug design. *para*-Aminosalicylic acid (PAS), remains a cornerstone therapy, in the treatment against *Mycobacterium tuberculosis*, owing to its high level of selectivity. Despite its high level of selectivity, PAS has been reassigned to treat drug-resistant strains of *M. tuberculosis* because it causes severe gastrointestinal (GI) distress that results in poor patient compliance. We have previously shown PAS inhibits the folate biosynthetic pathway specifically inhibiting dihydrofolate reductase^1,2^. In this study, we sought to determine the mechanistic basis of PAS selectivity and determined that PAS can be utilized in folate biosynthesis by other bacterial pathogens. The utilization of PAS ultimately led to the antagonism of key antibiotics, specifically the sulfonamides, used to prophylactically treat individuals with HIV-AIDS^3^. In addition, we found many bacteria in the GI tract could also utilize PAS to make a hydroxy-folate species which resulted in GI toxicity. Using sulfonamides as a tool to prevent PAS associated toxicity in the GI tract, we discovered that the sulfonamides antagonized the antimycobacterial activity of PAS. These findings indicate a new need for understanding the mechanisms of selective therapies and more important, that HIV-AIDS/*M. tuberculosis* co-infected individuals should avoid co-treatment of PAS and sulfonamides.

## Main text

Tuberculosis (TB) is the leading cause of death worldwide, particularly for individuals co-infected with the human immunodeficiency virus (HIV)^4^. Undoubtedly, a watershed moment in the fight against TB was the implementation of multi-drug therapy, which has attained cure rates of 85% for new cases of drug-sensitive TB^4^. Despite this dramatic achievement, combination therapy just for first-line antitubercular drugs requires 6 months of intensive treatment and is accompanied by severe side effects. The lengthy and multi-drug treatment is needed to combat the recalcitrant nature of *M. tuberculosis*, the causative agent of TB, which entails its complex cell wall structure, unusually slow metabolism, and propensity to acquire drug resistance. Over 70 years ago, the first synthetic antimicrobial for TB, *para*-Aminosalicylic acid (PAS), entered clinical use as a foundational cornerstone in multi-drug therapy used to treat *M. tuberculosis*. With nanomolar potency, PAS specifically targets the *M. tuberculosis* dihydrofolate reductase (FolA_*Mtb*_) and lacks activity against other microorganisms and mammalian cells^1,5–10^. Although highly selective against *M. tuberculosis* and listed as a World Health Organization (WHO) Model List of Essential Medicines, PAS has been reassigned to treat drug-resistant TB due to its association with severe gastrointestinal distress and challenge for patient adherence^4,11,12^. In this study, we sought to determine the mechanistic basis for PAS selectivity and toxicity. Understanding the mechanisms that govern the selectivity and toxicity of PAS will allow for the development of better tolerated and selective antitubercular agents.

Folate biosynthesis is conserved among all bacteria and is required for one carbon metabolism in the synthesis of DNA, RNA, and proteins. PAS is a structural analog of the folate biosynthesis precursor, *para*-aminobenzoic acid (PABA), and follows similar incorporation into the folate biosynthetic pathway (Extended Data Figure 1)^1,2,9,10,13^. PAS, like PABA, is ligated to 6-hydroxymethyl dihydropterin pyrophosphate by the enzyme dihydropteroate synthase (FolP) to produce the dihydropteroate analog, hydroxy-dihydropteroate (Extended Data Figure 1)^2,9,10^. Hydroxy-dihydropteroate is converted to the dihydrofolate (DHF) analog, hydroxy-DHF via DHF synthase (FolC) (Extended Data Figure 1)^2,9,10^. DHF is reduced by FolA to produce the folate, tetrahydrofolate. Although folate biosynthesis is a highly conserved pathway, it is unknown why hydroxy-DHF has activity against FolA_*Mtb*_. To better understand the selectivity of PAS, we sought to determine if PAS could be utilized in folate biosynthesis in other bacterial species.

Previous work has suggested that PAS has minimal activity against *Escherichia coli* and *Bacillus anthracis*^6–8,14^. Specifically, PAS rescued *E.coli* mutants that were PABA autotrophs^14^. We extended these previous studies to determine the minimum inhibitory concentration (MIC) of PAS required to inhibit growth against multiple bacteria including enteric commensal bacteria, Gram positive pathogens, Gram negative pathogens, non-pathogenic and non-tuberculosis mycobacteria. We found that all tested bacteria were resistant to PAS (Extended Data Table 1) suggesting that FolA in these bacteria could utilize hydroxy-DHF. We previously determined that hydroxy-DHF inhibited FolA in *M. tuberculosis* and in the present study we further interrogated if FolA was the principal target for hydroxy-DHF^1^. Since the non-pathogenic mycobacterium, *Mycobacterium smegmatis* (*Ms*), is resistant to PAS, we determined if PAS could be utilized in a *M. tuberculosis* Δ *pabB* strain expressing *in trans folAMtb* or *folA* from *M. smegmatis* (*folA_Ms_*). *M. tuberculosis* Δ *pabB* strain expressing *folA_Mtb_* grew only in the presence of PABA, not PAS (Extended Data Figure 2a)^15^. Interestingly, *M. tuberculosis* Δ *pabB* expressing *folA_Ms_* grew in the presence of both PAS and PABA (Extended Data Figure 2). Since both strains could not synthesize PABA, any growth in the presence of PAS suggests that FolA_*Ms*_ is capable of using hydroxy-DHF for one carbon metabolism. Thus, we conclude that FolA governs susceptibility and resistance to PAS. Next, we performed a similar experiment using *Escherichia coli*, which was also found to be resistant to PAS. We used an *E. coli* Δ*folA_Ec_* Δ*thyA* mutant expressing *in trans folA_Mtb_* and *folA_Ec_*. These strains cannot grow without exogenous supplementation of thymidine because the strain also lacks thymidylate synthase, a key enzyme in the thymidine synthesis pathway^16^. *E. coli* Δ*folA_Ec_* Δ*thyA* expressing *folA_Ec_* grew in the presence of thymidine and PAS. As expected, *E. coli* Δ*folA_Ec_* Δ*thyA* expressing *folA_Mtb_* grew only in the presence of thymidine alone and not in the presence of thymidine and PAS (Figure 1). Together these data indicate that FolA determines susceptibility to PAS.

**Figure 1.**
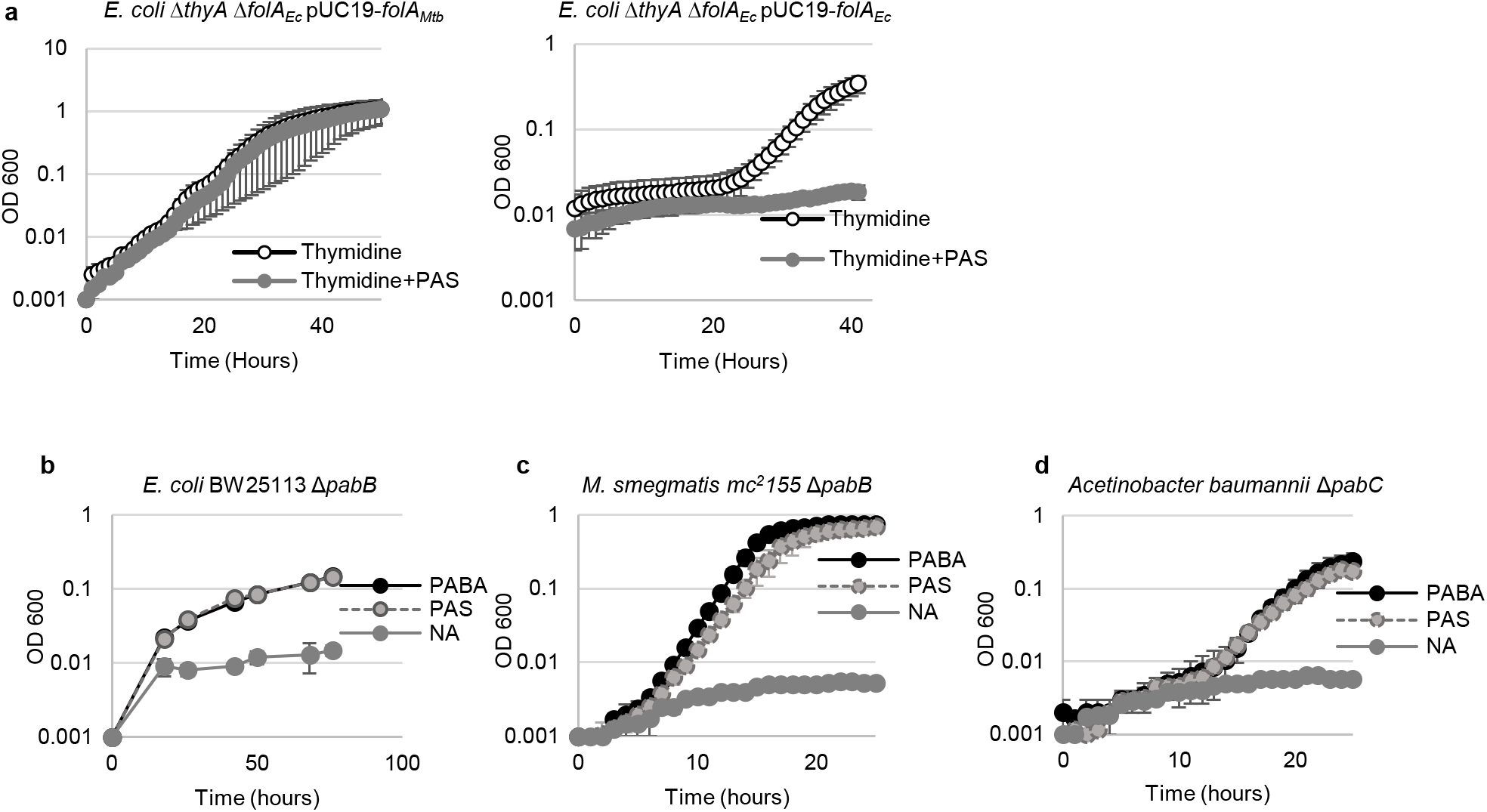
The mechanistic basis for PAS selectivity. *E. coli* BW25113 Δ*pabB* and *M. smegmatis mc^2^155* Δ*pabB* can utilize PAS in *lieu* of PABA for folate biosynthesis. **a,** *E. coli* Δ*thyA* Δ*folA_Ec_* pUC19-*folA_Ec_* can grow in the presence of thymidine and thymine+PAS (50μg/mL) in M9 minimal media. *E. coli* Δ*thyA* Δ*folA_Ec_* pUC19-*folA_Mtb_* can only grow in the presence of thymidine and not in the presence of thymine+PAS (50μg/mL) in M9 minimal media. **b,** *E. coli* BW25113 Δ*pabB*, **c,** *M. smegmatis mc^2^155* Δ*pabB*, and **d,** *Acetinobacter baumannii* Δ*pabC* were grown in minimal media the presence of 10μg/mL PABA or 10μg/mL. Data shown for growth curves are an average of three independent experiments error bars represent standard deviation.

We purified recombinant FolA from *E. coli* and *M. smegmatis* to determine if FolA from bacteria that are resistant to PAS could utilize hydroxy-DHF as a substrate. Both enzymes utilized DHF and hydroxy-DHF as substrates (FolA_*Ec*_ K_m DHF_ 2.1μM and K_m hydroxy-DHF_ 17.2μM, FolA_*Ms*_ K_m DHF_ 4.2 μM and K_m hydroxy-DHF_ 22.5 μM, Extended Data Figure 2). These data suggest that hydroxy-DHF can be utilized as a substrate for FolA in bacteria that are resistant to PAS. To determine if PAS was able to be utilized in one carbon metabolism, we used PABA auxotroph strains of *E. coli BW91120* Δ*pabB, M. smegmatis mc^2^155* Δ*pabB*, and *Acetinobacter baumannii ΔpabC*. When the strains were supplemented with exogenous PAS, the growth defects of all strains were chemically complemented, implying that PAS can be used *in lieu* of PABA in one carbon metabolism in bacteria that are resistant to PAS (Figure 1). Together these data suggest that PAS and, therefore, hydroxy-DHF are utilized as substrates for folate biosynthesis and, ultimately, one carbon metabolism.

It is intriguing how FolA_*Mtb*_ failed to use hydroxy-DHF as a substrate, since all bacterial FolA’s perform the same enzymatic function. To better understand this selectivity, we computationally docked DHF and hydroxy-DHF into crystal structures of FolA_*Mtb*_ and FolA_*Ec*_ (Extended Data Figure 2). The docking score was determined for each conformation of each enzyme-substrate complex and corresponds to the theoretical binding affinity, Kd. A more negative docking score in Glide correlates to a higher theoretical binding affinity of the enzyme for the substrate. In the FolA_*Mtb*_ hydroxy-DHF model, the hydroxyl group on hydroxy-DHF lies 2.4 Å away from Q30 and is able to form an additional hydrogen bond as compared to DHF (Extended Data Figure 3). This additional hydrogen bond would enable for hydroxy-DHF to be more tightly bound as seen by the more negative docking score. In the FolA_*Ec*_, the alpha helix adjacent to the binding site, residues 28-32, contains only hydrophobic amino acids that are unable to form any additional hydrogen bonds with hydroxy-DHF as compared to DHF. Evolutionary coupling was also used as a complementary approach to discover which residues were important in the selective binding of hydroxy-DHF. Evolutionary coupling predicted many residues to be conserved in FolA, however, only two residues were found to be coupled in the active site of FolA_*Mtb*_ I22 and Q30 (Extended Data Figure 3). Although the hydrophobic I22 position is conserved among FolA orthologs, Q30 is unique to FolA_*Mtb*_ and docking studies show that this residue plays a key role in the hydrogen bonding network of hydroxy-DHF (Extended Data Figure 3). The data suggests selectivity of PAS could be the result of an additional hydrogen bond with Q30 in FolA_*Mtb*_ which is absent in other FolAs resistant to PAS.

Since we demonstrated PAS can act *in lieu* of PABA in folate biosynthesis, we wanted to determine any resulting drug-drug interactions. Exogenous PABA can antagonize the activity of many FolP inhibitors such as sulfonamides and diaminodiphenyl sulfones^11,12,17,18^ (Figure 2 and Extended Data Figure 4). Unlike FolP inhibitors, exogenous PABA cannot antagonize the activity of trimethoprim (TMP), a FolA inhibitor^12,19^. HIV/AIDS-infected individuals are given a lifelong prophylactic combination therapy of SMX-TMP in addition to receiving anti-retroviral therapy^3^. We hypothesized that PAS also antagonizes the activity of sulfonamides and diaminodiphenyl sulfones but not TMP. Addition of exogenous PAS in *E. coli* antagonized the activity of the commonly prescribed sulfonamides including sulfamethoxazole (SMX), sulfanilamide, sulfathiazole, and the diaminodiphenyl sulfone, dapsone (Figure 2 and Extended Data Figure 4). Exogenous PAS had no interaction with TMP in *E. coli* (Figure 2 and Extended Data Figure 4). SMX and TMP are given in combination therapy and rarely given in monotherapy in the clinic. We tested if PAS could antagonize the activity of SMX and TMP given in a therapeutic 5:1 ratio. The minimum inhibitory concentration (MIC) of SMX in monotherapy is 0.2 μg/mL and the MIC of TMP in monotherapy is 0.5 μg/mL. The synergistic MIC of SMX and TMP is 0.194/0.0388 μg/mL, respectively (Figure 2 and Extended Data Figure 4). Surprisingly, PAS abrogated SMX and TMP synergy (Figure 2 and Extended Data Figure 4). To this end, once TMP exerts an MIC as found in monotherapy of 0.5 μg/mL, PAS can no longer antagonize the combined activity of SMX and TMP. This suggests that PAS can only antagonize the synergistic activity of SMX and TMP, resulting in an effect similar as treating with TMP alone. This data suggests treatment with PAS and SMX in patients co-infected with HIV/AIDS and *M. tuberculosis* may be contraindicated and should be reevaluated.

**Figure 2.**
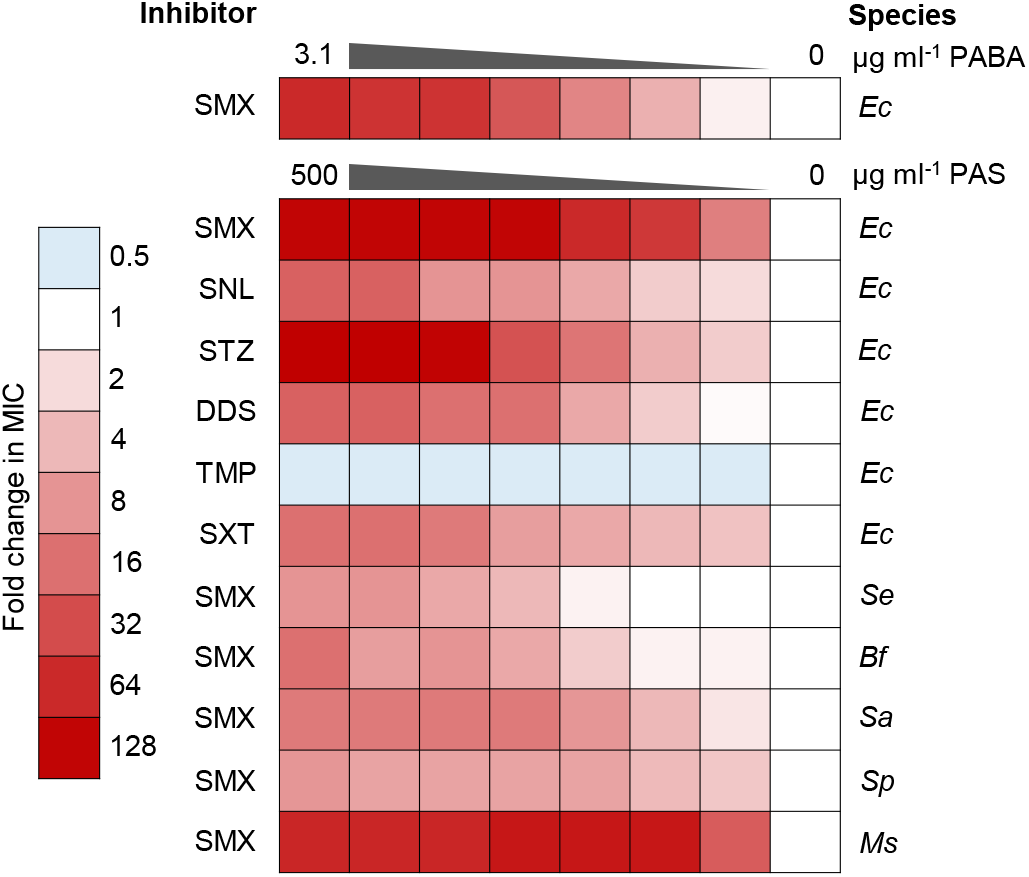
PAS can antagonize SMX activity in a variety of bacteria *in vitro*. Interactions between PAS and PABA, PAS and SMX, PAS and dapsone, PtePAS (PAS-pteroate) and SMX in M. tuberculosis H37Rv and H37Rv Δ*pabB* and various bacterial species. Data shown for growth curves are an average of three independent experiments. *Escherichia coli (Ec), Salmonella enterica* (*Se*), *Bacteroides fragilis (Bf), Staphyloccocus aureus (Sa), Streptococcus parasanguinis (Sp), Mycobacterium smegmatis* (*Ms*).

We also determined if exogenous PAS could antagonize SMX activity in a variety of other bacterial species. Exogenous PAS antagonized SMX activity against *M. smegmatis, Staphylococcus aureus, Streptococcus parasanguinis, Bacteroides fragilis*, and *Burkholderia cenocepacia*. PAS can be a substrate for folate biosynthesis precursor, and like PABA, can antagonize sulfonamides and sulfone activity in a variety of bacterial species (Figure 2 and Extended Data Figure 4). Together, these data represent a novel drug interaction between PAS and FolP inhibitors. The mechanistic basis for this antagonism relies on PAS susceptibility.

Humans lack many of the *de novo* enzymes required to make folates and must acquire folates from diet or the environment, specifically by the commensal microbiota^12^ Since we determined PAS can be utilized as a substrate for one-carbon metabolism, we hypothesized bacteria in the human gastrointestinal tract could produce a hydroxy-folate species. We reasoned that this hydroxy-folate species could result in toxicity as many patients discontinue PAS treatment because of the severe side effects, which largely encompasses gastrointestinal distress. In fact, we found that mice treated with PAS showed general signs of distress followed by a dose-dependent 63% mortality rate (Extended Data Figure 5) within 14 days of treatment. We reasoned enterocytes may utilize hydroxy-DHF or hydroxy-folate produced by colonic bacteria in response to PAS treatment as a source of folates. We found that purified human DHF reductase (DHFR) could use hydroxy-DHF as a suboptimal substrate (K_m_ 0.16 μM) compared to the native substrate, DHF (K_m_ 1.26 μM) (Extended Data Figure 5).

Enterocytes require a large concentration of folates for the maintenance of many cellular processes. Since human-DHFR could not utilize hydroxy-DHF as efficiently as DHF, enterocytes may not maintain normal cellular processes requiring folates. Conversely, the hydroxyl-group on hydroxy-DHF could be inhibitory in downstream metabolic processes. Cytotoxicity assays were performed using PAS, hydroxy-DHF and hydroxy-folate. PAS was not cytotoxic at physiologically relevant concentrations to HepG2 and Caco-2 cells (IC_50_ 1270 ± 9 μM and 1230 ± 6 μM, respectively) (Extended Data Table 2 and 3). Hydroxy-DHF was slightly cytotoxic in HepG2 and Caco-2 cells (IC_50_ 343 ± 4 μM and 370 ± 5 μM, respectively) (Extended Data Table 2 and 3). It is important to note that hydroxy-DHF is not stable in ambient oxygen; therefore, the full extent of its cytotoxicity in the anaerobic colon is unknown^1^. Interestingly, we found that hydroxy-folate, a reduced folate analog that is stable in oxygen, was cytotoxic in both HepG2 and Caco-2 cells (IC_50_ 35 ± 5 μM and 44 ± 5 μM, respectively) at similar concentrations to methotrexate (IC_50_ 7.2 ± 4.7 μM and 12 ± 6 μM, respectively), a known human-DHFR inhibitor (Extended Data Table 2 and 3)^12^. These data suggest that hydroxy-DHF/folate, and not PAS, are cytotoxic to human cells and may explain the mechanistic basis for PAS toxicity.

Since PAS toxicity was mediated by bioactivation of PAS into hydroxy-DHF or hydroxy-folate as a biproduct from the bacteria in the gastrointestinal tract, we hypothesized toxicity could be prevented by blocking PAS bioactivation. To test this hypothesis we co-treated mice with PAS and SMX, since SMX blocks FolP, the first enzyme required to convert PAS into hydroxy-dihydropteroate (Extended Data Figure 1). Groups of *M. tuberculosis* infected mice were treated with PBS (vehicle control), SMX (150 mg/kg), PAS (500 mg/kg or 750 mg/kg), or SMX and PAS (150 mg/kg and 750mg/kg, respectively) for 13 days via oral gavage. The mice in the vehicle control (n=6) and SMX treatment group (n=6) showed a 100% survival rate (Figure 3). Mice in the PAS (500 mg/kg) group (n=16) showed an 80% survival rate and mice in the PAS group (750 mg/kg) group (n=8) showed a 38% survival rate (Figure 3). The mice in the PAS-SMX (PAS 750 mg/kg and SMX 150 mg/kg) (n=8) co-treatment group showed an 87% survival rate (Figure 3). These observations suggest that SMX can antagonize PAS-mediated toxicity in mice by blocking the ability of bacteria in the gastrointestinal tract from converting PAS into hydroxy-DHF/folate.

**Figure 3.**
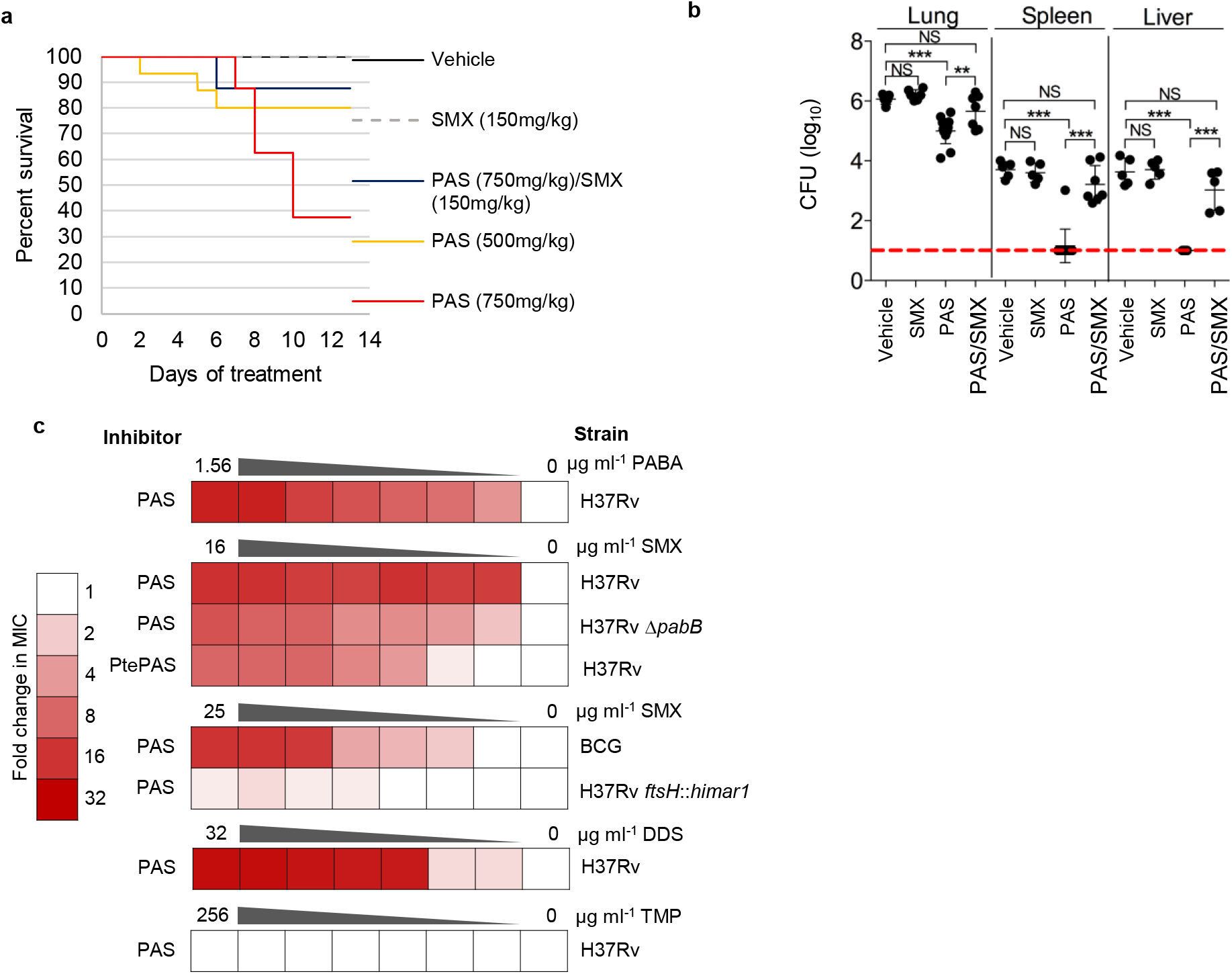
FolP inhibitors antagonize the anti-tubercular activity of PAS. **a,** Kaplan-Meier survival analysis of C57Bl6 mice aerosol infected with ~100 CFU *M. tuberculosis*. The infection was established for 1 week and the mice were treated for 13 days via daily oral gavage with vehicle control, SMX (150mg/kg), PAS (500mg/kg), and SMX/PAS (150mg/kg;500mg/kg) using an acute infection model. **b,** Bacterial burden at 14 days of treatment was enumerated in the lung, spleen, and liver in mice following treatment. The limit of detection (100 CFU) is marked by a red dotted line. **p<0.05, ***p<0.0001, NS (not significant) based on a students T-test. **c,** *In vitro i*nteractions between PAS and PABA, PAS and SMX, PAS and DDS, PtePAS and SMX in *M. tuberculosis* H37Rv and H37Rv Δ*pabB*. Data shown for *M. tuberculosis in vitro* antagonism assays are average of three independent experiments.

The bacterial burden was measured following treatment of the TB infected mice in the lung, spleen, and liver. The vehicle control and SMX treatment groups showed similar bacterial burdens in all organs (Figure 3). PAS treatment was able to reduce the burden of *M. tuberculosis* by 10-fold in the lungs of mice and virtually eliminated appearance of bacilli (below limit of detection) in spleen and liver (Figure 4). Co-treatment with SMX antagonized PAS antitubercular activity and resulted in an increase in bacterial burden in lung (~10-fold), liver (~1000-fold) and spleen (~1000-fold) compared to PAS treatment alone, that was statistically indistinguishable from the vehicle control and SMX treatment groups (Figure 3). These observations are consistent with previous work that has shown SMX antagonism of PAS *in vitro* (Figure 4)^10^. We extended this study to include dapsone, and found that dapsone can antagonize the antitubercular activity of PAS *in vitro* (Figure 4). Trimethoprim (TMP) had no interaction with PAS *in vitro* (Figure 4). Taken together, these data suggest that addition of a FolP inhibitor antagonizes the anti-mycobacterial activity of PAS *in vitro* and *in vivo*.

**Figure 4.**
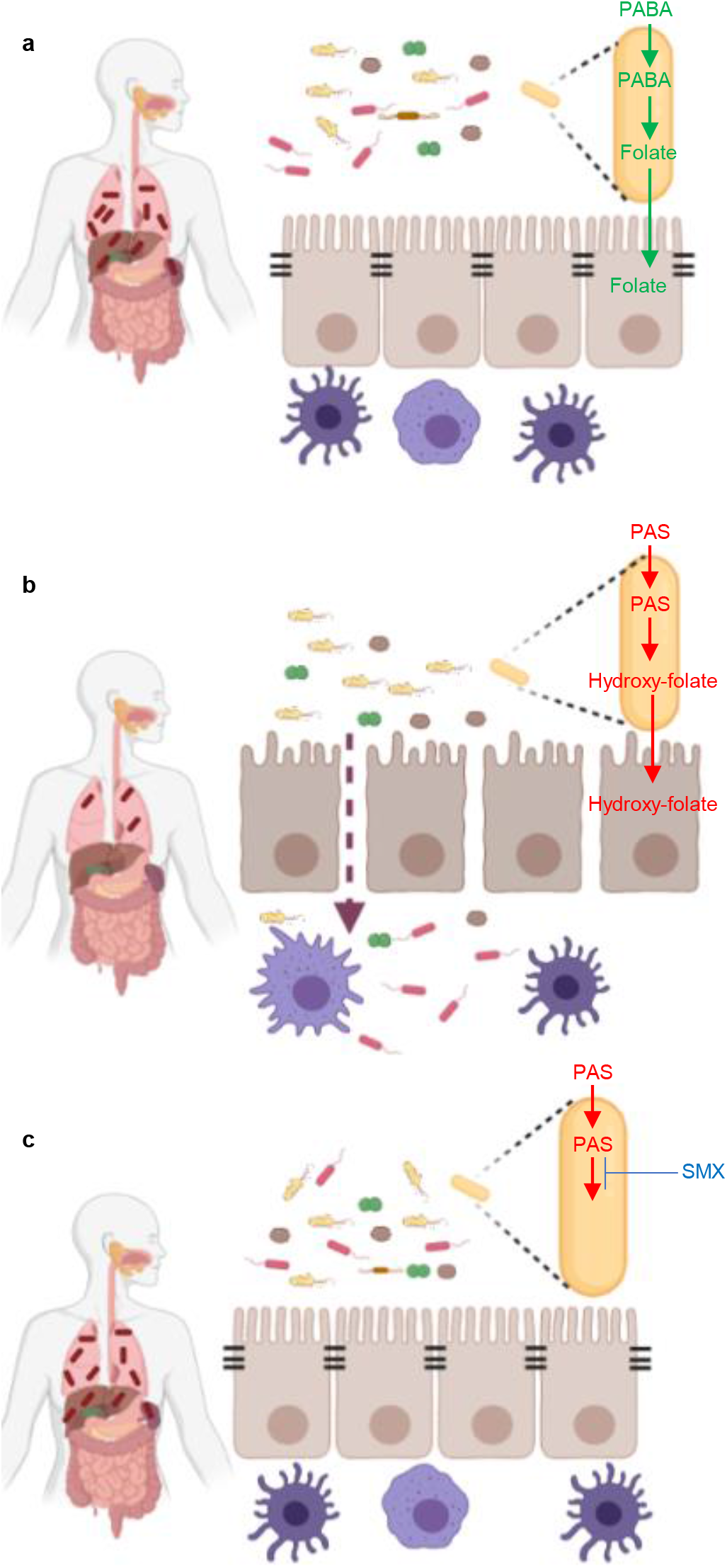
The systemic interactions between PAS and SMX in a *M. tuberculosis* infected individual. **a,** *M. tuberculosis* is able grow in the lungs and disseminate in into the liver and spleen without treatment. Commensal microbiota and intestinal epithelial cells unabated. **b,** *M. tuberculosis* replication is slowed during PAS treatment with some growth in the lungs. PAS results in intestinal epithelial cell death leading to the entry of commensal bacteria into the lumen. **c,** Co-treatment of SMX and PAS results in unrestricted growth of *M. tuberculosis* in the lungs, liver, and spleen. SMX antagonizes PAS toxicity in the gastrointestinal tract and intestinal epithelial cells and commensal bacteria are unabated and.

We hypothesized SMX antagonism is mediated by preventing PAS incorporation into the folate biosynthesis pathway. To test this hypothesis, we determined if SMX could antagonize hydroxy-pteroate, an oxidized form of hydroxy-dihydropteroate, to bypass PAS incorporation into hydroxy-dihydropteroate (Extended Data Figure 1)^20^. Interestingly, hydroxy-pteroate activity was antagonized by exogenous SMX (Figure 4 and Extended Data Figure 6). The data suggests that SMX antagonism of PAS is not occurring by preventing the incorporation of PAS to hydroxy-DHF. Exogenous PABA can also antagonize the activity of PAS (Figure 3 and Extended Data Figure 6)^10^. Since SMX treatment has previously been shown to increase PABA production in bacteria, we reasoned that SMX is causing an increase in PABA biosynthesis and, thereby, antagonism of PAS^17^. Previous work has shown that strains lacking *pabB* in *M. tuberculosis* are PABA auxotrophs and can only grow with exogenously added PABA^15^. We tested if SMX could antagonize PAS in *M. tuberculosis H37Rv Δ pabB* strain and found SMX could still antagonize PAS (Figure 3 and Extended Data Figure 6). This suggests that PABA biosynthesis partially mediates antagonism of PAS by SMX. Previous studies have shown that FolA inhibitors, such as TMP, can also modulate 6-hydroxymethyl dihyrodropterin pyrophosphate biosynthesis. Therefore we created a disruption in 6-hydroxymethyl dihyrodropterin pyrophosphate biosynthesis and found that *Mycobacterium bovis* BCG *ftsH::himar1* abolished SMX mediated PAS antagonism (Figure 4 and Extended Data Figure 6)^17^. These data suggest that SMX antagonism of PAS is mediated primarily through 6-hydroxymethyl dihyrodropterin pyrophosphate biosynthesis and also to a lesser extent PABA biosynthesis further corroborating that these two pathways are interconnected.

## Conclusion

This study describes the mechanistic basis for selectivity of one of the first antitubercular agents, PAS. Since its introduction into the clinic, PAS was noted for its precise selectivity for *M. tuberculosis*. Indeed, PAS exerted no deleterious effect against a variety of other bacterial species. We determined that PAS could be utilized as a substrate for folate metabolism in a variety of bacterial species except in *M. tuberculosis* and PAS susceptibility was restored with the integration of FolA_*Mtb*_. *M. tuberculosis* could utilize and grow in the presence of PAS with the addition of FolA_*Ms*_. We discovered that this selectivity mapped to key amino acid differences in FolA_*Mtb*_, the principle target of hydroxy-DHF, the active version of PAS. Furthermore, we uncovered this selectivity resulted in antagonism of SMX by PAS in a variety of bacterial species. SMX is used to prophylactically to prevent and treat opportunistic infections in patients with HIV/AIDS such as many of the bacteria tested in this study. Tuberculosis patients co-infected with HIV/AIDS are prescribed PAS and SMX treatment for prophylaxis. The data suggests that co-treatment with PAS may reduce SMX activity and negatively impact the health of TB patients co-infected with HIV (Figure 4). Furthermore, we also reasoned that this selectivity may be associated with PAS toxicity in humans. Human enterocytes cannot synthesize folates and rely on folates from diet or the GI microbiota. Indeed hydroxy-folate, an oxidized version of hydroxy-DHF, not PAS, was cytotoxic in HepG2 and Caco-2 cell lines. To prevent PAS associated toxicity, we reasoned co-treatment with SMX could prevent PAS from getting metabolized to hydroxy-DHF/hydroxy-folate and could alleviate toxicity. In a murine model, co-treatment of PAS and SMX prevented PAS associated toxicity; however, treatment with SMX resulted in increase in bacterial burden of mice infected with *M. tuberculosis* (Figure 4). The mechanistic basis for SMX mediated antagonism of PAS is due to an increase in pterin biosynthesis. The data from this study shows these two drugs, PAS and SMX should not be used in combination because both prevent the other from exerting the correct anti-microbial properties. This mutual antagonism is a direct result of PAS selectivity. Although antimicrobials that selectively inhibit an infectious agent are desirable, this study demonstrates that the mechanisms that govern selectivity need to be fully understood for future drug development. However, using the knowledge that PAS is selective for FolA_*Mtb*_ could allow for the development of novel inhibitors that fail to cause mutual antagonism.

## Acknowledgements

Pterin-PAS was a gift from Dr. Richard Lee and hydroxy-folate was a gift from Dr. Courtney Aldrich. We would like to thank Dr. Evan Krystofiak for helpful discussions. The authors would like to thank Shravika Talla and Abbey Hammes for their technical support. Research was supported by NIH grant R01 Al123146 awarded to A.D.B. SLK was funded through a University of Minnesota Doctoral Dissertation Fellowship. EAL was funded through a postdoctoral fellowship from the Ford Foundation and the National Academies of Science, Engineering, and Mathematics and NIH diversity supplement award (R01Al123146-03S1). MDH and AAB were funded through a University of Minnesota-Undergraduate Research Opportunity Program.

## Author contributions

SLK and ADB formulated the original hypothesis and designed the study. SLK performed the experiments and data analysis. EAL performed the *ex vivo* assays, assisted SLK with the mouse experiments, and analyzed the data. MDH and AAB assisted SLK in performing the bacteriology assays. WM and BF performed the molecular docking studies. SLK, EAL, WM, BF, and ADB wrote the manuscript. All authors commented on the manuscript, data, and conclusions.

## Methods

### Bacterial strains, media, and growth conditions

All information regarding primers, plasmids, and bacterial can be found in Extended Data Table 4.

*M. tuberculosis* strains and *Mycobacterium bovis* BCG were grown at 37 °C in either 7H9 broth (Difco) supplemented with 0.2% (vol/vol) glycerol (Fisher Scientific), 10% (vol/vol) oleic acid-albumin-dextrose-catalase (OADC) (Difco), and 0.05% (vol/vol) tyloxapol (Sigma-Aldrich) or 7H10 agar (Difco) supplemented with 0.2% glycerol and OADC.

All antibiotics were added when appropriate to final concentrations of 50μg/mL for kanamycin, and 150 μg/mL for hygromycin. To eliminate PABA contamination, glassware was baked for a minimum of one hour at 180 °C. PAS, PABA, trimethoprim (TMP), dapsone (DDS), and methotrexate (MTX) were purchased from Sigma and were dissolved in 100% DMSO (Sigma). 2’-Hydroxy-7,8-dihydrofolate was synthesized as described previously^1^. Pterin-PAS and hydroxy-folate were dissolved in 100% DMSO (Sigma).

### *Biochemical characterization of FolA*_Mtb_, *FolA*_Ec_ and *FolA*_Ms_

Cloning of *M. tuberculosis folA* was performed as described previously^1^.

*E. coli* BW25113 *folA* was amplified using primers in Supplementary Table (X).

The resulting DNA was cut with the restriction enzymes *NdeI* and *Bam*HI and ligated into an already digested pET28b(+) using the same restriction enzymes. Expression and purification of FolA_*Ec*_ was similarly performed as previously described^21^. Briefly, sequence-verified pET28b(+):*folA_Mtb_* was transformed into competent *E. coli* BL21 (DE3) cells. E. coli BL21 pET28b(+)-*folA_Ec_* was inoculated into LB and grown overnight at 37 °C. The cells were diluted 1:1000 into fresh LB (1 L) and were grown until mid-exponential phase (OD_600_ 0.4–0.6) at 37 °C and 1 mM IPTG was added to induce protein expression at 37 °C for 4 h. The cells were collected by centrifugation at 5,000 rpm at 4 °C. The pellet was resuspended in 10 mL of lysis buffer (50 mM NaH2PO4, 300 mM NaCl, and 10 mM imidazole (pH 8.0)) containing 10 mg chicken egg white lysozyme was added and incubated on ice for 30 min. The insoluble fraction was removed by centrifugation at 11,000 rpm at 4 °C for 45 min. The supernatant was applied to a 1 mL of Ni-NTA Agarose (Qiagen) equilibrated with lysis buffer. The column was washed with 10 mL of wash buffer (50 mM NaH2PO4, 300 mM NaCl, and 20 mM imidazole (pH 8.0)). The protein eluted and collected in 5-1mL aliquots of elution buffer (50 mM NaH2PO4, 300 mM NaCl, and 250 mM imidazole (GoldBio) (pH 8.0)). Fractions containing pure FolA_*Ec*_ (>90% as judged by an SDS-PAGE gel) were pooled, concentrated (Millipore) into storage buffer (25 mM Tris buffer (pH 7.5) with 10% glycerol and 1 mM DTT) to 10 mg/mL. The protein was stored at −80 °C.

FolA_*Ms*_ was purified similarly as the purification of FolA_*Mtb*_. Briefly, *M. smegmatis folA_Ms_* was amplified using *M. smegmatis* genomic DNA by PCR. The resulting DNA was cut with the restriction enzymes *NdeI* and *BamH1* and ligated into an already digested pET28b(+) using the same restriction enzymes. Briefly, sequence-verified pET28b(+): *folA_Ms_* was transformed into competent *E. coli* BL21 (DE3) cells. E. coli BL21 pET28b(+): *folA_Mtb_* was inoculated into LB and grown overnight at 37 °C. The cells were diluted 1:1000 into fresh LB (1 L) and were grown until mid-exponential phase (optical density at 600 nm 0.4-0.6) at 37 °C and 1 mM IPTG was added to induce protein expression at 37 °C for 4 h. The cells were collected by centrifugation at 5,000 rpm (Beckman Coulter, Avanti JXN-30) at 4 °C. The pellet was resuspended in 10 mL of lysis buffer (20 mM triethanolamine (TEA), 50 mM KCl, pH 7) and was ultrasonicated (Branson Sonifier 450) three times using 20 sec burst (4 °C) and 20 sec cooling. 10 mg chicken egg white lysozyme (MP Biomedicals, LLC) was added and incubated on ice for 30 min. The insoluble fraction was removed by centrifugation (Beckman Coulter, Avanti JXN-30) at 11,000 rpm at 4 °C for 45 min. The supernatant was applied to a 1 mL of Ni-NTA Agarose (Qiagen) equilibrated with lysis buffer. The column was washed with 40 mL of wash buffer (20 mM TEA, 50 mM KCl, 50 mM imidazole (GoldBio), pH 7). The protein was eluted with 5 mL of elution buffer (20 mM TEA, 50 mM KCl, 500 mM imidazole (GoldBio), pH 7). Fractions containing pure FolA_*Ms*_ (>90% as judged by an SDS-PAGE gel) were pooled, concentrated (Millipore) into storage buffer (20 mM potassium phosphate, 50 mM KCl, pH 7.0) to 5 mg/ml. The protein was stored at −80 °C.

### Biochemical utilization of DHF and hydroxy-DHF

All enzymatic assays were performed in flat bottom 96 well plates (Corning), with 200 μL reaction volume, and measured in a BioTek Synergy H1 spectrophotometer at 25 °C. FolA_*Ec*_ enzymatic assays were performed using 5 nM enzyme in MTEN buffer [50 mM 2-morpholinoethanesulfonic acid, 25 mM tris(hydroxymethyl)aminomethane, 25 mM ethanolamine, and 100 mM NaCl (pH 7.0)] containing 1 mM DTT and 0.01% (vol/vol) Triton-X 100. The enzyme was preincubated with 67 μM of NADPH at room temperature for 5 min. The reaction was initiated with varying concentrations of dihydrofolate or hydroxy-dihydrofolate. The decrease in absorbance corresponding to NADPH oxidation was monitored at 340 nm every 10 sec for 10 min. Kinetic measurements for FolA_*Ms*_ were performed identically as FolA_*Ec*_ except the reaction was performed in 20 mM potassium phosphate, 50 mM KCl, pH 7.0 with 0.01% (vol/vol) Triton-X 100. The K_m_ were determined from 4 independent experiments performed in biological triplicate and analyzed using GraphPad Prism software.

### Construction of PABA auxotrophic strains

*E. coli ΔpabB* and *A. baumannii ΔpabC* strains were constructed as previously described^22^ *M. smegmatis pabB* gene was replaced with a hygromycin resistance and *sacB* cassette using the specialized transduction method^23^ Briefly, ~1,000 bp regions upstream and downstream of *pabB* were amplified via PCR, digested with *Van91I* and ligated into previously digested p0004S. The resulting plasmid was sequenced to verify amplicons. The resulting plasmid was digested with *PacI* and ligated into previously digested phAE159. The resulting temperature sensitive phage was propagated at 30 °C in *M. smegmatis* to high titer and used to transduce *M. smegmatis*. The resulting transductants were plated on 10 μg/mL PABA and hygromycin and incubated at 37 °C. The deletion was verified by PCR. H37Rv Δ*pabB* was created identically to *M. smegmatis ΔpabB* except using primers found in Table 2.1 and the transduced H37Rv was plated on 1 μg/mL PABA and hygromycin.

### PABA auxotroph growth curves

*E. coli ΔpabB* and *A. baumannii ΔpabC* were grown to mid-exponential phase (OD_600_ 0.4–0.6) in LB medium and washed three times with PABA-free M9 medium. The cells were subcultured in PABA-free M9 medium to OD_600_ 0.001 in the presence of 10 μg/mL PABA, 10 μg/mL PAS, or no addition, in technical triplicate, in round bottom 96-well plates (Corning). The cells were incubated at 37 °C, with 200 rpm shaking, and OD_600_ were read every hour for 24 hrs. *M. smegmatis* Δ*pabB* was grown in PABA-free supplemented 7H9 containing 10 μg/mL PABA to mid-exponential phase (OD_600_ 0.4-0.6) and washed three times with PABA-free 7H9. The cells were subcultured in PABA-free 7H9 to an OD_600_ of 0.001 in the presence of 10 μg/mL PABA, 10 μg/mL PAS, or no addition, in technical triplicate, in round bottom 96-well plates (Corning). The cells were incubated at 37 °C without shaking. Absorbance (OD_600_) was read every 6 and 18 hours (Spectronic Genesys 5S). All growth curves were performed in biological triplicate.

### E. coli ΔthyA ΔfolA *containing pUC19 constructs*

*E. coli folA* was amplified using primers in Table 2.1 using PCR. The resulting DNA was cut with the restriction enzymes *Bam*HI and *XmaI* and ligated into an already digested pUC19. The resulting plasmid was sequence verified and electroporated into *E. coli* Δ*thyA* Δ*folA. M. tuberculosis folA* was amplified from a codon optimized G Block (Invitrogen) using primers in Table 2.1 for PCR. The resulting DNA was cut with restriction enzymes *Bam*HI and *Eco*RI and ligated into an already digested pUC19. The resulting plasmid was sequence verified and electroporated into *E. coli* Δ*thyA* Δ*folA*.

### *H37Rv* ΔpabB *containing pUMN002 constructs*

*M. smegmatis folA* and *M. tuberculosis folA* were amplified using primers in Table 2.1 for PCR. The resulting amplicons were cut with the restriction enzymes *HindIII* and *Eco*RI and ligated into an already digested pUMN002. The resulting plasmids was sequence verified and electroporated into H37Rv Δ*pabB*. The strains were plated on 7H10 containing 1 μg/mL PABA, hygromycin, and kanamycin.

### folA *swap growth curves*

*E. coli* Δ*thyA* Δ*folA* containing either pUC19-*folA_Ec_ or pUC19-folA_Mtb_* was grown in M9 medium supplemented with 200 μg/mL thymine (Sigma) and 50 μM IPTG to mid-exponential phase (OD_600_ 0.4-0.6) and washed three times in M9 medium. *E. coli* containing pUC19-*folA_Ec_* was subcultured to OD_600_ 0.001 and *E. coli* containing pUC19-*folA_Mtb_* was subcultured to OD_600_ 0.01, in M9 supplemented with 200 μg/mL thymine (Sigma) and 50 μM IPTG, either alone or with 50 μg/mL PAS, in round bottom 96-well plates (Corning), in technical triplicate. The cells were incubated at 37 °C with shaking at 200rpm. Absorbance (OD_600_) was read every hour for 50 hours. All growth curves were performed in biological triplicate.

H37Rv Δ*pabB* containing pUMN002, pUMN002-*folA_Mtb_* or pUMN002-*folA_Ms_* was grown with 10 ng/mL PABA to mid-exponential phase and was subcultured to OD_600_ 0.01 with 10 ng/mL PABA. The strains were grown to mid-exponential phase, subcultured to OD_600_ 0.1, and streaked on PABA-free 7H10 plates containing 5 μg/mL PAS or 5 μg/mL PABA. The plates were incubated for 3 weeks at 37 °C.

### Computational docking

Dihydrofolate and hydroxy-DHF were each docked into the crystal structures of analogous FolA_*Ec*_ and FolA_*Mtb*_. Docking was performed using the Schrödinger Maestro suite [Schrödinger, LLC, New York, NY, 2018]. The X-ray crystal structures of *E. coli* DHFR with methotrexate and NADPH (PDB 4P66) and *Mtb* DHFR with methotrexate and NADPH (PDB 1DF7) were retrieved from the RCSB PDB^24–26^ The PDB structures were prepared for docking studies using the Maestro Protein Preparation Wizard to assign bond orders, create disulfide bonds, fill-in gaps in the protein structure, and add in hydrogens. Waters greater than 5 Å away from a ligand or amino acid hetero group were removed and then an energy minimization was completed using the OPLS3 force field. The Maestro Receptor Grid Generation module was used to define a 25 × 25 × 25 Å grid centered on the methotrexate position in both structures. Dihydrofolate and hydroxy-DHF ligand structures for docking were prepared by adaptation from the methotrexate structure of 4P66 using the Maestro 3D Build Module. Docking was performed using the Maestro Glide module-with extra precision and flexible ligand sampling, but no additional constraints. 10 initial poses were generated for each molecule and subjected to a post-docking minimization using an OPLS3 force field. The resulting poses were ranked according to their Glide score, which is an approximation of binding energy.

### Evolutionary coupling

The evolutionary coupling between pairs of residues in FolA_*Mtb*_ was determined using EVcouplings (http://www.EVfold.org)^27,28^. The amino acid sequence used was UniProt ID <u>P9WNX1</u> and the PDB 1DG8 was used for the structural comparisons^25^. All other default parameters were used. The majority of coupling pairs are required for structural folding of the protein and map outside of the active site. The only coupling pair found within the active site was Q28 and I20.

### Determination of minimum inhibitory concentrations of antifolates for bacterial strains

Minimum inhibitory concentration (MIC_90_) is defined as the minimum concentration of antimicrobial agent required to inhibit ≥90% of growth compared to a no drug control. MIC_50_ is defined as the minimum concentration of drug required to inhibit ≥50% of growth compared to a no drug control. Growth was assessed spectrophotometrically (OD_600_) (BioTek Synergy H1) or visually, when noted.

*M. tuberculosis* was grown to mid-exponential phase and subcultured to OD_600_ 0.01 in inkwell bottles. The MIC_90_ of PAS was determined using log_2_ serial dilutions. The MIC_90_ was determined after 14 days of incubation. The MIC_90_ and checkerboards of PAS and SMX for *E. coli* and *S. aureus* were performed as previously described^17^ using log_2_ serial dilutions. *M. smegmatis* was grown to mid-exponential phase and subcultured to OD_600_ 0.001 in a round bottom 96-well plate (Corning). The MIC_90_ and checkerboards for PAS and SMX were performed using log_2_ serial dilutions and the MIC_90_ was determined after 3 days of static incubation. *A. baumannii, S. enterica, S. maltophilia*, and *B. cenocepacia* were grown in LB to mid-exponential phase and washed 3 times with M9 medium. The cells were diluted to OD_600_ 0.001 in M9 medium in round bottom 96-well plates (Corning). The MIC and checkerboards for PAS and SMX were performed using log_2_ serial dilutions and the MIC (or MIC for *B. cenocepacia*) was determined visually after 24 hrs (or 3 days for *B. cenocepacia)* of static incubation. *B. fragilis* was grown in BHIS to mid-exponential phase and washed three times with AMMGluc. The cells were diluted to OD_600_ 0.001 in AMMGluc in round bottom 96-well plates (Corning). The MIC and checkerboards for PAS and SMX were performed using log_2_ serial dilutions and the MIC was determined visually after 3 days of static incubation. *S. parasanguinis* was grown in Mueller-Hinton broth to mid-exponential phase and washed three times with Iso-Sensitest broth. The cells were diluted to OD_600_ 0.001 in Iso-Sensitest broth in flat bottom 96-well plates (Corning). The MIC_90_ and checkerboards for PAS and SMX was performed using log_2_ serially dilutions and the MIC_50_ was determined visually after 24 hrs of static incubation. All MICs were performed in biological triplicate.

### Cell Lines

Hep-G2 cell (ATCC HB-8065) and Caco-2 cell (ATCC HTB-37) lines were purchased from the American Type Culture Collection (ATCC; Manassas, VA). Hep-G2 and Caco-2 cell lines were maintained in Minimal Essential Medium (MEM; Gibco, Waltham, MA) and Dulbecco’s Modified Eagle Medium (DMEM; Gibco), respectively. Media were supplemented with 10% (Hep-G2) or 20% (Caco-2) fetal bovine serum (FBS) and 1% penicillin/streptomycin (pen/strep) solution. Both cell lines were incubated at 37 °C in a humidified chamber containing 5% CO2. Medium was refreshed every 2 days. Once 70% confluency was achieved, cell lines were washed thrice using Dulbecco’s phosphate buffered saline (D-PBS) without calcium and magnesium (Gibco) and harvested with TrypLE™ express enzyme (1X; Gibco). Detached cells were subsequently used in cytotoxicity assays.

### Cytotoxicity assays

Unless otherwise noted, cells were incubated at 37 °C in a humidified chamber containing 5% CO2. Hep-G2 and Caco-2 cells were seeded separately in tissue culture treated, flat-bottom 96 well plates at a density of 5.0 x 10^4^ cells per well in antibiotic free media. The final volume per well was 100 μL. After cells were adhered to the well substrate overnight, cells were washed thrice with D-PBS and incubated with individual drugs and metabolites, methotrexate (MTX; 0-4,000 μM), *para*-aminosalicylic acid (PAS; 0-4,000 μM), hydroxy-folate (0-500 μM), and hydroxyl-dihydrofolate (0-500 μM) in a two-fold serial dilution using appropriate culture media. Cells were treated for up to 72 h and media containing appropriate drugs were replaced every 24 h. DMSO vehicle control was included for every time point. Cell survival after drug exposure was determined using previously established methods^29^. Briefly, 200 μL of freshly made 3-(4,5-dimethylthiazol-2-yl)-2,5-diphenyltetrazolium bromide (MTT; 1 mg/mL; Sigma-Aldrich, St. Louis, MO) in serum-free, phenol red-free MEM or DMEM was added to each well and incubated for 3 h. MTT solution was removed and formazan crystals were dissolved in 200 μL of isopropanol. Formazan dye was quantified at 570 nm using the Synergy H1 microtiter plate reader (BioTek; Winooski, VT). Absorbance was normalized for background at OD650. Cell survival was calculated as the percentage absorbance of sample relative to no vehicle control. IC_50_ (half maximal inhibitory concentrations) values for each drug at all time points tested were calculated using GraphPad Prism (San Diego, CA) statistical software. All treatments were conducted in technical triplicate for each time point. All experiments were repeated thrice.

### Assessing PAS toxicity in mice

Seven week old C57BL/6 mice were obtained from Jackson Laboratory (Bar Harbor, ME). Female mice (5 per treatment) were orally gavaged every day for two weeks with a vehicle control of phosphate buffered saline (PBS) (pH 7.2) or 750 mg/kg *para*-aminosalicylic acid. Before the mice were gavaged, the cannula was submerged in a 10% (weight/volume) sucrose (Fisher) solution. Fecal pellets (3-6) were collected from individual mice prior to initiation of treatment, after two weeks of treatment, and after a two week recovery following treatment. Mice were sacrificed and the small and large intestines were harvested and washed in PBS.

### Construction and purification of DHFR_Human_

Human DHFR isoform 1 cDNA was codon optimized and purchased as a gene block (Invitrogen) containing 5’ *Nde*I and 3’ *Bam*HI cut sites. The gene block was digested with *Nde*I and *Bam*HI and ligated into an already digested pET28b(+) (Novagen) using the same restriction enzymes. Expression and purification of DHFRHuman was performed as previously described^30^. Briefly, sequence-verified pET28b(+):*DHFR_Human_* was used to transform competent *E. coli* BL21 (DE3) cells. *E. coli* BL21 pET28b(+):*DHFR_Human_* was inoculated into Lysogeny Broth (LB) and grown overnight at 37 °C. The cells were diluted 1:1000 into fresh LB (4 L) and were grown until mid-exponential phase (optical density at 600 nm (OD_600_) 0.4–0.6) at 37 °C. Next, 1 mM isopropyl β-D-1-thiogalactopyranoside (IPTG) (GoldBio) was added to induce protein expression at 37 °C for 4 h. The cells were collected by centrifugation at 5,000 rpm (Beckman Coulter, Avanti JXN-30) at 4 °C. The pellet was resuspended in 10 mL of lysis buffer (100 mM K_2_PO_4_ (pH 8.0) and 5 mM imidazole) and disrupted by ultrasonication (Branson Sonifier 450) three times using 20 sec burst and 20 sec cooling (4 °C). 10 mg chicken egg white lysozyme (MP Biomedicals, LLC) was added and incubated on ice for 30 min. The insoluble fraction was removed by centrifugation (Beckman Coulter, Avanti JXN-30) at 11,000 rpm at 4 °C for 45 min. The supernatant was applied to 1 ml Ni-NTA Agarose (Qiagen) equilibrated with lysis buffer. DHFR_Human_ was eluted using a step-wise gradient of 10 mL wash buffer containing increasing concentrations of imidazole (10 mM, 15 mM and 20 mM). DHFRHuman was eluted with 5 mL of elution buffer (100 mM K_2_PO_4_ (pH 7.5) and 50 mM imidazole). Fractions containing pure DHFRHuman (>90% as judged by running samples on an SDS-PAGE gel) were pooled and using an ultra-centrifugal filter concentrated (Millipore) in storage buffer (50 mM KPO_4_, 5 mM β-mercaptoethanol, pH 7.3) to 2.5 mg/ml. The protein was stored at 4 °C.

### Biochemical utilization of DHF and hydroxy-DHF

All enzymatic assays were performed in flat bottom 96 well plates (Corning), with 200 μl reaction volume, and measured in a BioTek Synergy H1 spectrophotometer at 25 °C. The enzymatic reactions were performed as previously described^31^. Enzyme assays were performed using 5 nM enzyme in 50 mM KPO_4_, pH 7.3, 5 mM β-mercaptoethanol and 0.01% (vol/vol) Triton-X 100. The enzyme was preincubated with 67 μM of NADPH at room temperature for 5 min. The reaction was initiated with varying concentrations of DHF or hydroxy-DHF. The decrease in absorbance corresponding to NADPH oxidation was monitored at 340 nm every 10 sec for 10 min. The K_m_ was determined from 4 independent experiments performed in biological triplicate and analyzed using GraphPad Prism software.

### *Bacterial enumeration of* M. tuberculosis *infected mice during antitubercular oral gavage treatment*

Seven week old C57BL/6 mice were obtained from Jackson Laboratory (Bar Harbor, ME). Mice were infected with ~100 CFU of *M. tuberculosis* H37Rv using an inhalation exposure system (GlasCol) as previously described^32^. The infection was established for 1 week. Following the 1 week incubation period, mice were gavaged daily for 13 days with vehicle (PBS), SMX (150 mg/kg), PAS (500 mg/kg or 750 mg/kg), and a combination of SMX (150 mg/kg) and PAS (750 mg/mg). Before gavage, the cannula was submerged in 10% (wt/vol) sucrose (Fisher) solution. Following each daily treatment mice were fed peanut butter. The peanut butter (PB2 powdered peanut butter) (Amazon) was suspended in 50:50 (weight/volume) of sterile water. Infected mice were euthanized by CO2 overdose. Bacterial CFU were enumerated by plating serially diluted lung, spleen, and liver homogenates on complete Middlebrook 7H10 agar containing 100 μg/ml cycloheximide. The CFU’s were enumerated after 3 to 4 weeks of incubation at 37 °C. All animal protocols were reviewed and approved by the University of Minnesota Institutional Animal Care and Use Committee and were conducted in accordance with recommendations in the National Institutes of Health Guide for the Care and Use of Laboratory Animals. The protocols, personnel and animals used were approved and monitored by the Institutional Animal Care and Use Committee.

### M. tuberculosis *antagonism assays*

All assays were performed in round bottom 96-well plates (Corning) except where noted. Minimum inhibitory concentration (MIC_90_) is defined as the concentration to inhibit 90% of growth compared to a no drug control. Growth was assessed spectrophotometrically (OD_600_) (BioTek Synergy H1) or visually, when noted. All assays were performed in biological triplicate.

*M. tuberculosis* H37Rv was grown to mid-exponential phase and subcultured to OD_600_ 0.001 in 96 round bottom plates (Corning). The interactions between PABA and PAS, PAS and SMX, PAS and DDS, and pterin-PAS and SMX were evaluated using log_2_ serial dilutions. The MIC was determined visually after 14 days of static incubation at 37 °C. The interactions between TMP and PAS, using log_2_ serial dilutions, was performed in *M. tuberculosis* H37Rv grown to mid-exponential phase and subcultured to OD_600_ 0.01 in inkwell bottles at 37 °C, with shaking. The MIC_90_ of TMP and PAS was measured spectrophotometrically (GENESYS 20, Thermo Fisher) after 14 days.

*M. tuberculosis* H37Rv Δ*pabB* was constructed as described in Chapter 2. *M. tuberculosis* H37Rv Δ*pabB* was grown in 7H9 medium containing 1 μg/mL of PABA to mid-exponential phase and subcultured to OD_600_ 0.001 in 7H9 media containing 100ng/mL of PABA in round bottom 96-well plates (Corning). The interactions between PAS and SMX was performed using log_2_ serial dilutions. The MIC was determined visually after 14 days of static incubation at 37 °C.

*M. bovis* BCG and *M. bovis* BCG *ftsH::himar1* was grown in 7H9 medium to mid-exponential phase and subcultured to OD_600_ 0.001 in 7H9 media in round bottom 96-well plates (Corning). The interactions between PAS and SMX was performed using log_2_ serial dilutions. The MIC_90_ was determined in a BioTek Synergy H1 spectrophotometer after 14 days of static incubation at 37 °C.

### Statistical analysis

Number of mice required to produce results with statistical significance were determined by a power calculation. A sample size of n≥4 was used detect a 10-fold (1-log10) difference in CFU between groups, assuming standard deviations of 35-40% of sample mean, with a type 1 error rate (α) of 0.05% to achieve a 90% power^33^ A student’s unpaired *t* test (two tailed) was used for comparison between vehicle and treatment groups. *p*-values were calculated using GraphPad Prism 5.0 software (GraphPad Software, Inc.). *p* ≤ 0.05 was considered significant.

