## Extended data figures for "Mechanism of selectivity reveals novel antifolate drug interactions": Extended data figures 20200513.pdf

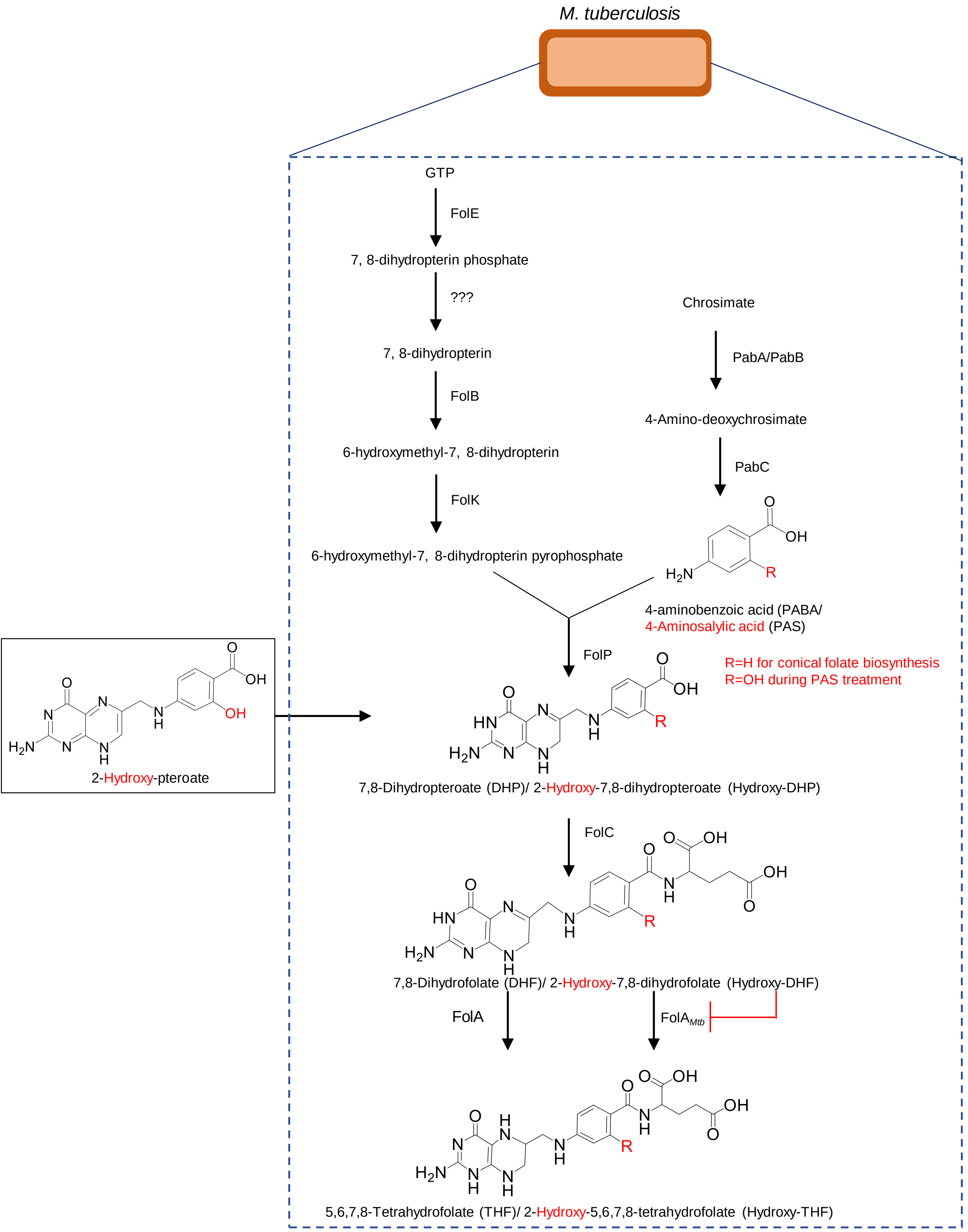


Extended Data Figure 1. Folate biosynthesis and PAS activation in bacteria. PAS is converted by the folate biosynthesis to hydroxy-DHF that inhibits $FolA_{Mtb}$ not FolA in other bacteria.

Extended Data Table 1. PAS Inhibitory concentrations against various bacterial species

| Organism | PAS MIC (µg/mL) |
|---|---|
| *Mycobacterium tuberculosis*[a] | 0.3 |
| *Mycobacterium smegmatis* | >500 |
| *Mycobacterium abscessus* | >500 |
| *Escherichia coli* | >500 |
| *Acetinobacter baumannii* | >500 |
| *Bacteroides fragilis* | 250 |
| *Burkholderia cenocepacia* | >500 |
| *Salmonella enterica* | >500 |
| *Stenotrophomonas maltophilia* | >500 |
| *Staphyloccocus aureus* | >500 |
| *Streptococcus parasanguinis* | 250 |

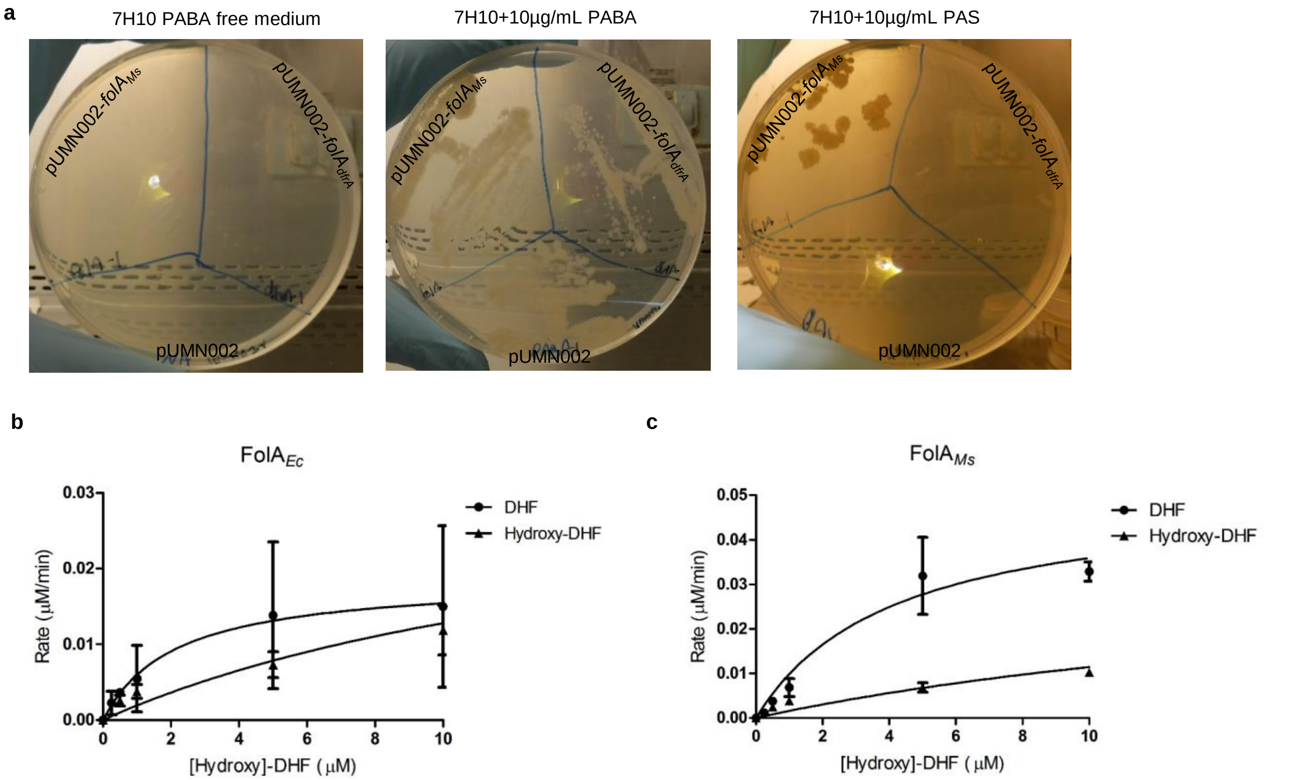


Extended Data Figure 2. FolA utilization of DHF or hydroxy-DHF in *E. coli*, *M. smegmatis*, and *M. tuberculosis*. **a,** H37Rv Δ*pabB* pumn002 can only grow in the presence of exogenous PABA (5µg/mL). H37Rv Δ*pabB* pumn002-*folA*$_{Ms}$ can grow in the presence of PABA (5µg/mL) and PAS (5µg/mL). Data shown represents three independent plating experiments. Dihydrofolate or hydroxy-dihydrofolate utilization was measured in the presence of dihydrofolate or hydroxy-dihydrofolate using purified recombinant dihydrofolate reductase from **b,** *E. coli* and **c,** *M. smegmatis*. The data represents the average error of the mean performed in technical triplicate with a minimum of four biological replicates.

**a**

FolA$_{Mtb}$ DHF

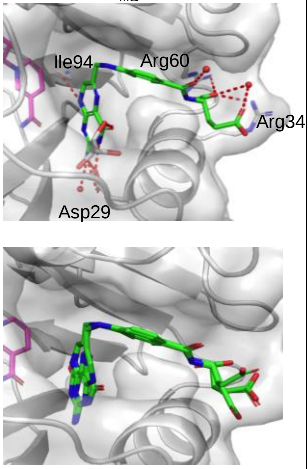


| Pose | FolA$_{Mtb}$ Docking Score (DHF) |
|---|---|
| 1 | -11.639 |
| 2 | -11.537 |
| 3 | -11.403 |
| 4 | -11.163 |
| 5 | -11.020 |

**b**

FolA$_{Mtb}$ Hydroxy-DHF

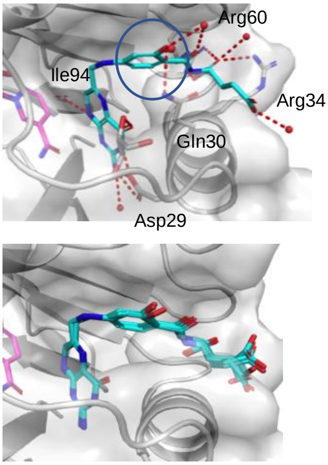


| Pose | FolA$_{Mtb}$ Docking Score (hydroxy-DHF) |
|---|---|
| 1 | -12.591 |
| 2 | -12.410 |
| 3 | -12.017 |
| 4 | -11.935 |
| 5 | -11.929 |

**c**

FolA$_{Ec}$ DHF

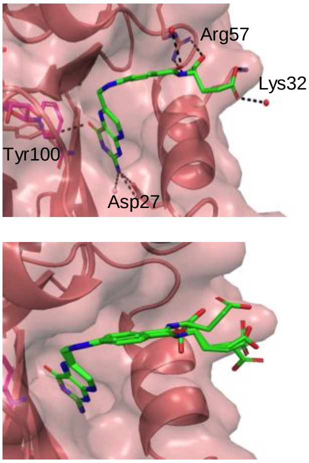


| Pose | FolA$_{Ec}$ Docking Score (DHF) |
|---|---|
| 1 | -9.440 |
| 2 | -9.370 |
| 3 | -9.322 |
| 4 | -9.028 |

**d**

FolA$_{Ec}$ Hydroxy-DHF

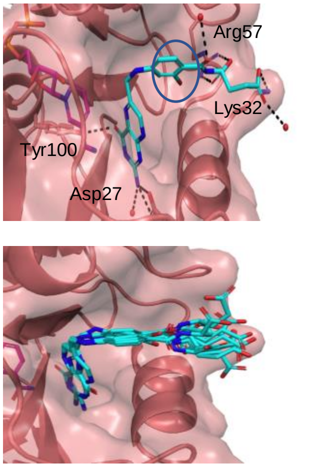


| Pose | FolA$_{Ec}$ Docking Score (hydroxy-DHF) |
|---|---|
| 1 | -9.177 |
| 2 | -8.948 |
| 3 | -8.598 |
| 4 | -8.483 |

**e**

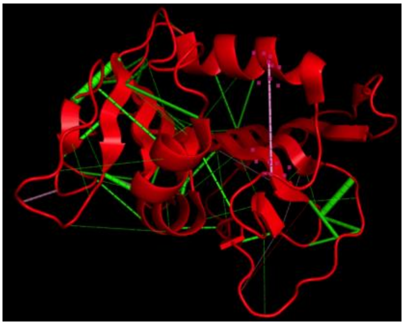

**f**

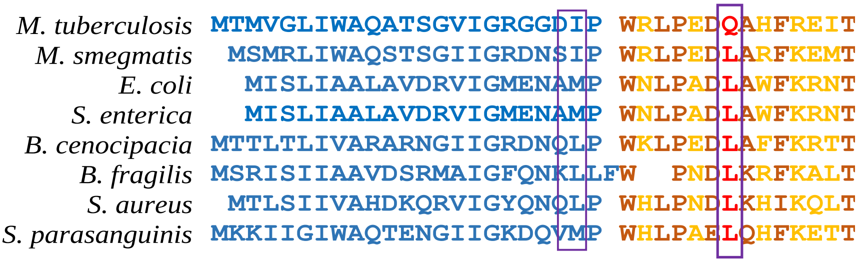


Extended Data Figure 3. Molecular docking and evolutionary coupling predict a hydrogen bond in FolA$_{Mtb}$. Previously crystallized FolA$_{Mtb}$ (PDB:1DF7) or FolA$_{Ec}$ (PDB 4P66) was used to dock a,c) DHF or b,d)hydroxy-DHF. Dashed lines represent key hydrogen bonds formed during the docking simulation. Red dots represent water molecules found in the crystal structures of the molecules used for docking. Circle indicated hydrogen bond formation present in FolA$_{Mtb}$ bound to hydroxy-DHF that is absent in FolA$_{Ec}$ bound to hydroxy-DHF. Molecular docking scores and positions of a,c) DHF and b,d) hydroxy-DHF in FolA$_{Mtb}$ and FolA$_{Ec}$, respectively. e) Evolutionary coupling was performed on FolA$_{Mtb}$, although multiple residues were considered coupled (green) the only residues in the active site found to be coupled were Q30 and I22. f) Amino acid sequence alignment of the Met20 loop shows conservation of I22 amino acid among FolA (box). Q30 is only found in is the only residue not conserved in the amino acid sequence comparison (box).

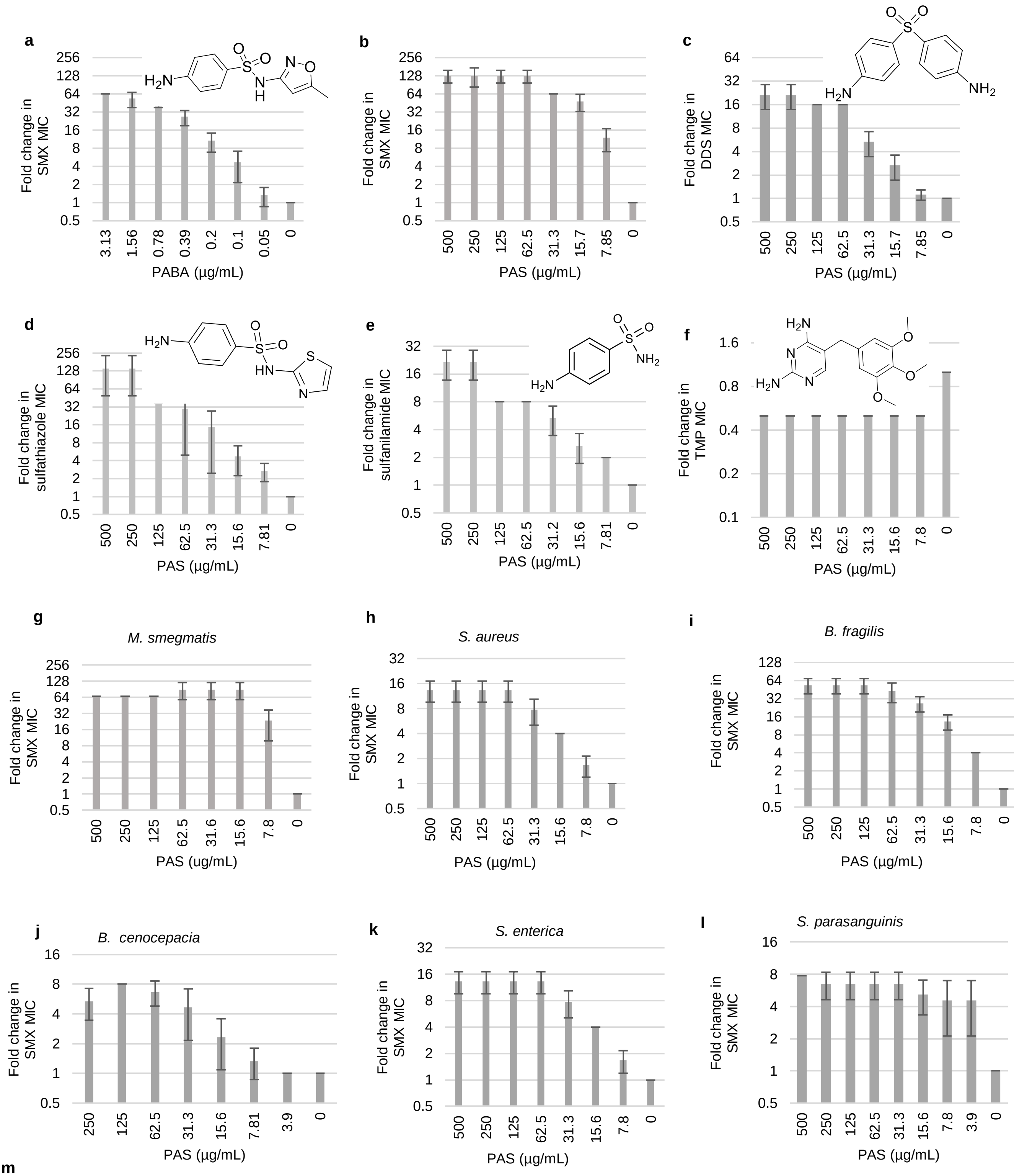


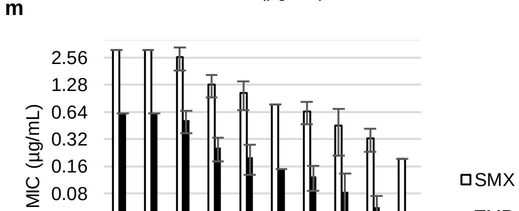


Extended Data Figure 3. PAS, like PABA, can antagonize antifolates various bacterial species. *E. coli* were grown in minimal M9 medium in the presence of varying concentrations of a) PABA and sulfamethoxazole (SMX). *E. coli* were grown in minimal M9 medium in the presence of varying concentrations of PAS and either b) sulfamethoxazole (SMX), c) dapsone (DDS), d) sulfathiazole (STZ). e) sulfanilamide (SNL), f) trimethoprim (TMP) Checkerboard assays were performed using PAS and SMX in various bacterial species. h) *M. smegmatis,* i) *S. aureus*, j) *B. fragilis,* k) *B. cenocepacia,* l) *S. enterica,* , m) *S. parasanguinis* were grown in minimal media, or isosentist broth for *S. parasanguinis*. m) PAS can only partially antagonize a clinical combination of SMX [illegible]

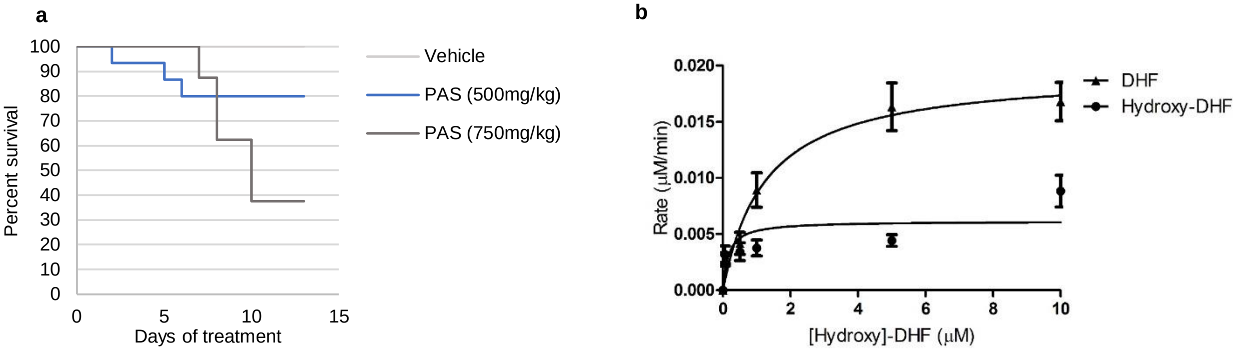


Extended Data Figure 5. The effect of PAS on mammalian DHFR. a) Specific pathogen free C57Bl/6 mice were treated via oral gavage with either a vehicle control (PBS) or 750 mg/kg PAS. The mice were treated every day for 14 days. Mice were assessed daily and examined for signs of distress including shortness of breath, withdrawal from littermates, and lethargy. Any mouse that exhibited these signs was euthanized. b) DHF and hydroxy-DHF can be utilized as substrates for $DHFR_{Human}$. DHF or hydroxy-DHF utilization was measured in the presence of DHF or hydroxy-DHF using purified recombinant DHFR. The experiment was performed in technical triplicate four times. The error bars represent standard error of the mean.

Extended Data Table 2. Cytotoxicity assays, in HepG2 cells, of MTX, PAS, hydroxy-DHF, and hydroxy-folate.

| HepG2 | 24hr | 48hr | 72hr |
| --- | --- | --- | --- |
| | $IC_{50}$(μM) | | |
| Methotrexate | 822.5 | 583.6 | 10.03 |
| PAS | 1706 | 1468 | 1468 |
| Hydroxy-DHF | 415.9 | 346 | 341.3 |
| Hydroxy-folate | --- | 163.8 | 39.98 |

Extended Data Table 3. Cytotoxicity assays, in Caco-2 cells, of MTX, PAS, hydroxy-DHF, and hydroxy-folate.

| Caco2 | 24hr | 48hr | 72hr |
| --- | --- | --- | --- |
| | $IC_{50}$(μM) | | |
| Methotrexate | 822.5 | 583.6 | 10.03 |
| PAS | 1706 | 1468 | 1468 |
| Hydroxy-DHF | 415.9 | 346 | 341.3 |
| Hydroxy-folate | --- | 163.8 | 39.98 |

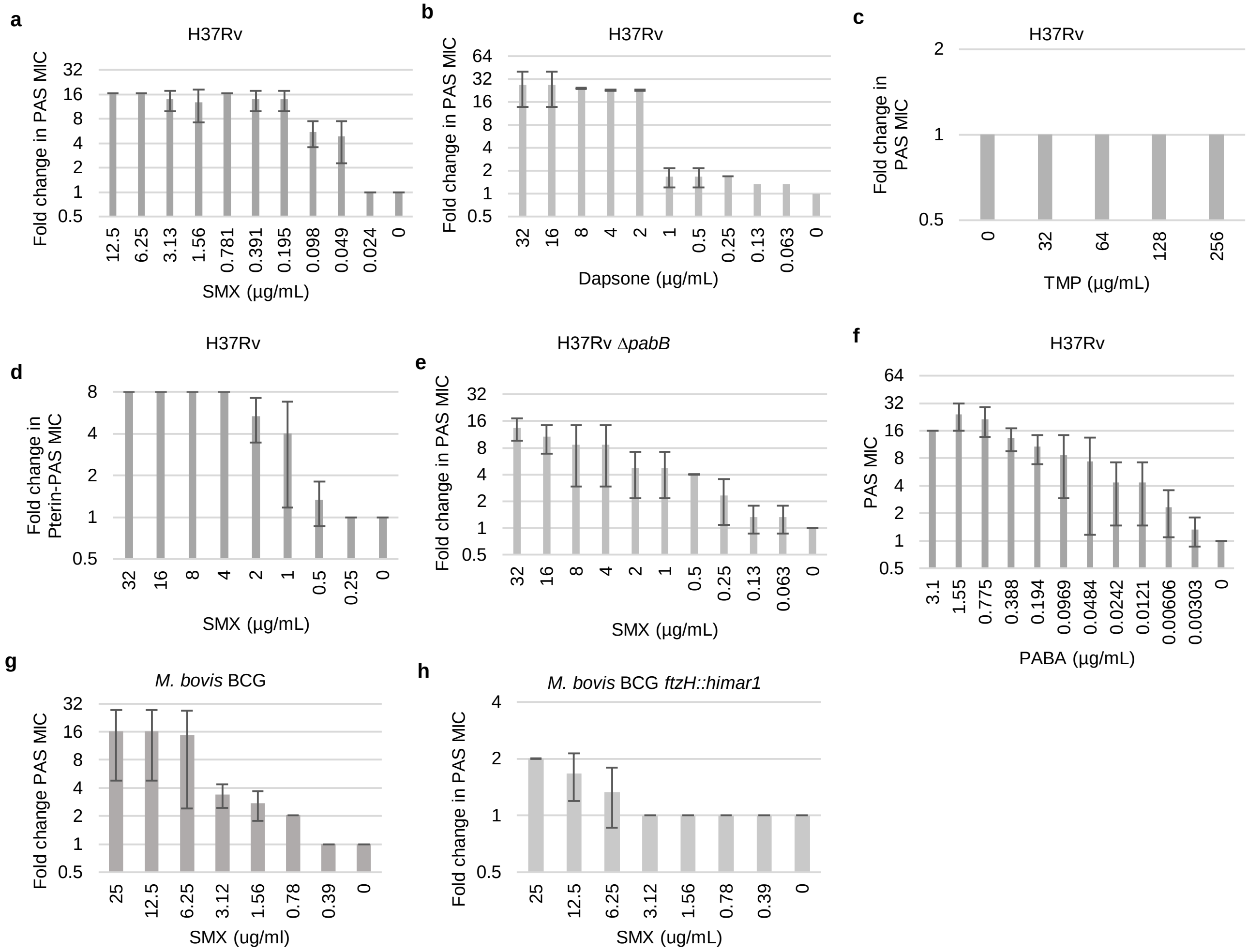


Extended Data Figure 6. FolP inhibitors antagonize the anti-tubercular activity of PAS. *in vitro* Interactions between PAS and PABA, PAS and SMX, PAS and DDS, PtePAS and SMX in *M. tuberculosis* H37Rv and H37Rv Δ*pabB*. *in vitro* Interactions between PAS and SMX in b) *M. bovis* BCG and c) *M. bovis* BCG *ftsH*::*himar*1. Strains were grown in 7H9 with appropriate supplementations at 37 ºC for 2 weeks. The data represents the minimum concentration of drug required to inhibit growth in varying concentrations. Data shown represent average of three independent experiments. Error bars represent standard deviation. Data shown represent average of three independent experiments..

Extended Data Table 4. Oligonucleotides, plasmids, and strains used in this study

| Primer name | Primer sequence[1] | Source | Restriction enzyme |
|---|---|---|---|
| pet28b-folA-Ec for | TTTTTTCATATGATGATCAGTCTG | This work | *Nde*1 |
| pet28b-folA-Ec rev | TTTTTTGGATCCTTACCGCCGC | This work | *Bam*HI |
| pabB_Ms LL | TTTTTTTTCCATAAATTGGTTCTGGTTCAAGGGACGCAACCGCG | This work | *Van*91I |
| pabB_Ms LR | TTTTTTTTCCATTTCTTGGGCCATCAGCGTGGCAGATTGCTAGC | This work | *Van*91I |
| pabB_Ms RL | TTTTTTTTCCATAGATTGGAGTTCGCGCTTGGACGGCGCGCCCG | This work | *Van*91I |
| pabB_Ms RR | TTTTTTTTCCATCTTTTGGCCCAGGTTCAAGAACCGGGTGATCG | This work | *Van*91I |
| pUC19-folA-Ec-for | TTTTTTGGATCCATGATCAGTCTGATTGCGGCGTT | This work | *Bam*HI |
| pUC19-folA-Ec-rev | TTTTTTCCCGGGTTACCGCCGCTCCAGAATCTC | This work | *Xma*1 |
| pUC19-folA-Mtb-for | TTTTTTGGATCCATGGTCGGCTTAATC | This work | *Bam*HI |
| pUC19-folA-Mtb-rev | TTTTTTGAATTCTCAACTACGATGGTAC | This work | *Eco*RI |
| pabB_Mtb LL | TTTTTTTTCCATAAATTGGCTCGCAAACTCGCGTCGTAGG | 15 | *Van*91I |
| pabB_Mtb LR | TTTTTTTTCCATTTCTTGGGCACCGGACAGGCTCTCATAC | 15 | *Van*91I |
| pabB-Mtb RL | TTTTTTTTCCATAGATTGGAGTGTGGCACCTGGTGTCCAC | 15 | *Van*91I |
| pabB-Mtb RR | TTTTTTTTCCATCTTTTGGACTCCAGCGCGTTAACCGCAA | 15 | *Van*91I |
| pUMN002-folA-Ms-for | TTTTTTAAGCTTATGATGAGACTGATCTGGGCCCA | This work | *Hind*III |
| pUMN002-folA-Ms-rev | TTTTTTGAATTCTCAGCGCCGGTAGCTGCAGA | This work | *Eco*RI |
| pUMN002-folA-Mtb-for | TTTTTTAAGCTTATGGTGGGGCTGATCTGGGCT | This work | *Hind*III |
| pUMN002-folA-Mtb-rev | TTTTTTGAATTCTCATGAGCGGTGGTAGCT | This work | *Eco*R |

| Plasmid | Relevant characteristics | Source |
|---|---|---|
| pET28b+ | Bacterial high copy plasmid for protein overexpression containing C- and N-terminal 6X histidine tags inducible with lacI, Kan$^r$ | Novagen |
| pET28b-folA-Mtb | pET28b+ containing folA$_{Mtb}$ with an N-terminal 6x-histidine tag, Kan$^r$ | 1 |
| pET28b-folA-Ec | pET28b+ containing folA$_{Ec}$ with an N-terminal 6x-histidine tag, Kan$^r$ | This work |
| pET28b-folA-Ms | pET28b+ containing folA$_{Ms}$ with an N-terminal 6x-histidine tag, Kan$^r$ | This work |
| p0004S | Plasmid used for allelic exchange, Hygro$^r$, sacB | 34 |
| phAE159 | Phasmid used for specialized transduction, Pen$^r$ | 23 |
| pUC19 | E. coli complementation plasmid, inducible with lacI, Pen$^r$ | Lab stock |
| pUC19-folA-Ec | pUC19 containing folA$_{Ec}$, Pen$^r$ | This work |
| pUC19-folA-Mtb | pUC19 containing folA$_{Mtb}$ that has been codon optimized for expression in E. coli, Pen$^r$ | This work |
| pUMN002 | Mycobacterial low copy number constitutively replicating plasmid, Kan$^r$ | Lab stock |
| pUMN002- folA$_{Ms}$ | pUMN002 containing folA$_{Ms}$, Kan$^r$ | This work |
| pUMN002-folA$_{Mtb}$ | pUMN002 containing folA$_{Mtb}$, Kan$^r$ | This work |
| pET28b-*DHFR*-human | pET28b+ containing *DHFR*$_{Human}$ with an N-terminal 6x-histidine tag, Kan$^r$ | This work |

| Strain Name | Relevant characteristics | Source |
|---|---|---|
| E. coli | | |
| DH5α | Cloning strain | Lab stock |
| BL21 (DE3) | Protein purification strain | 1 |
| pET28b-folA-Mtb | BL21 containing pET28b-folA-Mtb for overexpression of FolA$_{Mtb}$ | This work |
| pET28b-folA-Ec | BL21 containing pET28b-folA-Ec for overexpression of FolA$_{Ec}$ | This work |
| pET28b-folA-Ms | BL21 containing pET28b-folA-Ms for overexpression of FolA$_{Ms}$ | This work |
| BW25113 | Wild-type lab stock | Lab stock |
| ΔpabB | BW25113 pabB::kan, Kan$^r$ | Keio collection 22 |
| ΔthyA ΔfolA$_{Ec}$ | LH18, ΔthyA ΔfolA$_{Ec}$::kan, Kanr | 16 |
